# Neuroanatomical correlates of food addiction and obesity in the general population

**DOI:** 10.1101/411504

**Authors:** F. Beyer, I. García-García, M. Heinrich, M. Scholz, ML Schroeter, J. Sacher, T. Luck, S.G. Riedel-Heller, M. Stumvoll, A. Villringer, A.V. Witte

**Affiliations:** Department of Neurology, Max Planck Institute for Human Cognitive and Brain Sciences, Leipzig; Montreal Neurological Institute, McGill University, Montreal; EGG-Lab, Department of Neurology, Max Planck Institute for Human Cognitive and Brain Sciences, Leipzig; Institute for Medical Informatics, Statistics and Epidemiology, University of Leipzig, Leipzig; Department of Cognitive Neurology, University Hospital, Leipzig; Department of Neurology, Max Planck Institute for Human Cognitive and Brain Sciences, Leipzig; EGG-Lab, Max Planck Institute for Human Cognitive and Brain Sciences, Leipzig; Institute of Social Medicine, Occupational Health and Public Health (ISAP); Clinic for Endocrinology and Nephrology, University Hospital, Leipzig; Department of Neurology, Max Planck Institute for Human Cognitive and Brain Sciences, Leipzig Department of Cognitive Neurology, University Hospital, Leipzig

**Keywords:** Food addiction, Obesity, Body Mass Index, Gray Matter, Neuroimaging, Eating, Cohort Studies

## Abstract

The food addiction model suggests neurobiological similarities between substance-related and addictive disorders and obesity. While structural brain differences have been consistently reported in these conditions, little is known about the neuroanatomical correlates of food addiction. We therefore assessed whether food addiction, assessed with the Yale Food Addiction Scale (YFAS), related to obesity, personality and brain structure in a large population-based sample (n=625; 20-59 years old, 45% women). A higher YFAS symptom score correlated with obesity and disinhibited eating. In a whole-brain analysis, YFAS symptom score was not associated with cortical thickness nor subcortical gray matter volumes. Higher body mass index (BMI) correlated with reduced thickness of (pre)frontal, temporal and occipital cortex. Bayes factor analysis suggested that BMI and - to a smaller extent - YFAS symptom score contributed independently to right lateral orbitofrontal cortex thickness. Our study shows that food addiction is not associated with neuroanatomical differences in a large population-based sample, and does not account for the major part of obesity-associated gray matter alterations. Yet, food addiction might explain additional variance in orbitofrontal cortex, a hub area of the reward network. Longitudinal studies implementing both anatomical and functional MRI could further disentangle the neural mechanisms of addictive eating behaviors.

## Main text

The food addiction model provides a theoretical framework that explains the development of obesity based on observed similarities between substance addictions and overeating behavior (Randolph, 1956; Gearhardt et al., 2009). Essentially, this model proposes that some individuals exhibit eating patterns, elicited by certain types of food, that resemble addictive behaviors with regard to loss of control over eating and continued food consumption despite harmful consequences (Volkow et al., 2013b; Volkow et al., 2013a; Hebebrand et al., 2014; Schulte et al., 2015) but see (Ziauddeen et al., 2012).

Evidence in support of this hypothesis has been found in patients with eating disorders (Gearhardt et al., 2014) but also in epidemiological cohorts (Pedram et al., 2013; Flint et al., 2014; Pursey et al., 2016) where self-reported food addiction correlated with higher body mass index (BMI) and other measures of obesity.

Previous neuroimaging studies have highlighted the putative role of the prefrontal cortex and dopamine-dependent fronto-striatal circuits in dysfunctional self-regulation which could underlie both substance-related and addictive disorders and pathological eating behavior (Fineberg et al., 2010; García-García et al., 2014; Voon et al., 2015; García-García et al., 2017). For example, positron emission tomography studies reported lower dopamine D2/D3 striatal receptor binding both in participants with substance-related and addictive disorders (Volkow et al., 2008) and in morbidly obese individuals (Wang et al., 2001). Along these lines, participants with food addiction showed altered reward-related brain activity in the dorsolateral and orbitofrontal cortex and in the caudate nucleus in response to anticipated receipt of (rewarding) food (Gearhardt et al., 2011).

Regarding structural variation in addictive disorders, several studies reported lower grey matter volume (GMV) in the inferior and superior frontal cortex, orbitofrontal cortex and medial occipital gyrus that discriminated stimulant-dependent individuals from non-dependent controls (Durazzo et al., 2011; Fortier et al., 2011; Ersche et al., 2012; Moreno-López et al., 2012; Pennington et al., 2015). Lower GMV in middle and orbitofrontal areas also had a prognostic value in predicting alcohol dependence relapse after 12 months (Durazzo et al., 2011).

Similarly, obesity has been consistently associated with reduced GMV and lower cortical thickness in (pre)frontal cortex, the temporal lobe and bilateral cerebellum (Hassenstab et al., 2012; Marqués-Iturria et al., 2013; Kharabian Masouleh et al., 2016; García-García et al., 2018).

Several studies in middle-to aged populations raised the hypothesis that the observed gray matter volume loss is mainly a consequence of adverse metabolic factors related to obesity (Gustafson et al., 2004; Janowitz et al., 2015; Willette and Kapogiannis, 2015; Shahrzad Kharabian et al., 2017). Low-grade inflammation and dysregulated glucose metabolism, for example, may trigger damage to nervous tissue and contribute to accelerated brain aging and the increased risk for dementia associated with obesity (Beydoun et al., 2008; Baumgart et al., 2015; Ronan et al., 2016). Another line of research suggests that altered gray matter structure in brain regions that govern aspects of eating behavior may represent a risk factor for weight gain and obesity (Yokum et al., 2011; Hassenstab et al., 2012; Smucny et al., 2012; Opel et al., 2017).

Taken together, according to the food addiction model, addictive-like eating behavior is one of multiple factors that contribute to obesity. Akin neurobiological mechanisms might underlie both substance-related and addictive disorders and obesity (Volkow et al., 2008; Volkow et al., 2013b; García-García et al., 2014) and possibly account for impulsive and compulsive behaviors in these disorders (Michaud et al., 2017). Both addictive disorder and obesity are associated with alterations in gray matter structure, however, structural variation related to food addiction remains largely unexplored.

Here, we aimed to determine the relation of food addiction with obesity and brain morphology in a large population-based cohort. We hypothesized that food addiction was associated with structural alterations in fronto-striatal brain areas. To investigate the role of food addiction within the multiple factors involved in obesity (Hruby and Hu, 2015), we tested whether food addiction predicted different measures of obesity, including abdominal MRI-based body fat, as well as measures of eating behavior and general personality features (Vainik et al., 2013; Gerlach et al., 2014). We additionally assessed obesity-related differences in brain structure and compared the evidence for the involvement of BMI and food addiction in selected regions of interest using a Bayesian statistical approach.

## Methods

### Participants

All participants were enrolled in the “Health Study for the Leipzig Research Centre for Civilization Diseases” (LIFE-Adult) study (Loeffler et al., 2015). We selected adult participants from the neuroimaging cohort (n=2637) with available head MRI and without history of neurological or psychiatric disease such as stroke, cancer, epilepsy, multiple sclerosis and Parkinson’s disease. We restricted the age range to younger and middle-aged participants (20 to 59 years old) to avoid overestimation of aging-related effects in the brain among participants older than 60 years (Fjell et al., 2012; Storsve et al., 2014). Included participants did not take central nervous system agents and had complete data for the primary covariates Yale Food Addiction Scale (YFAS) and body mass index (BMI). Due to potential links between food addiction and depression (Davis et al., 2011), participants were excluded if they reported major depressive disorder in the last 12 months or scored > 21 in the CES-D (Center for Epidemiological Studies-Depression) (Radloff, 1977). In total, 625 participants met inclusion criteria, and were included into the main analysis (Figure 1). Out of these, n = 444 underwent abdominal MRI and n > 423 completed further questionnaires assessing eating behavior and personality.

**Figure 1.**
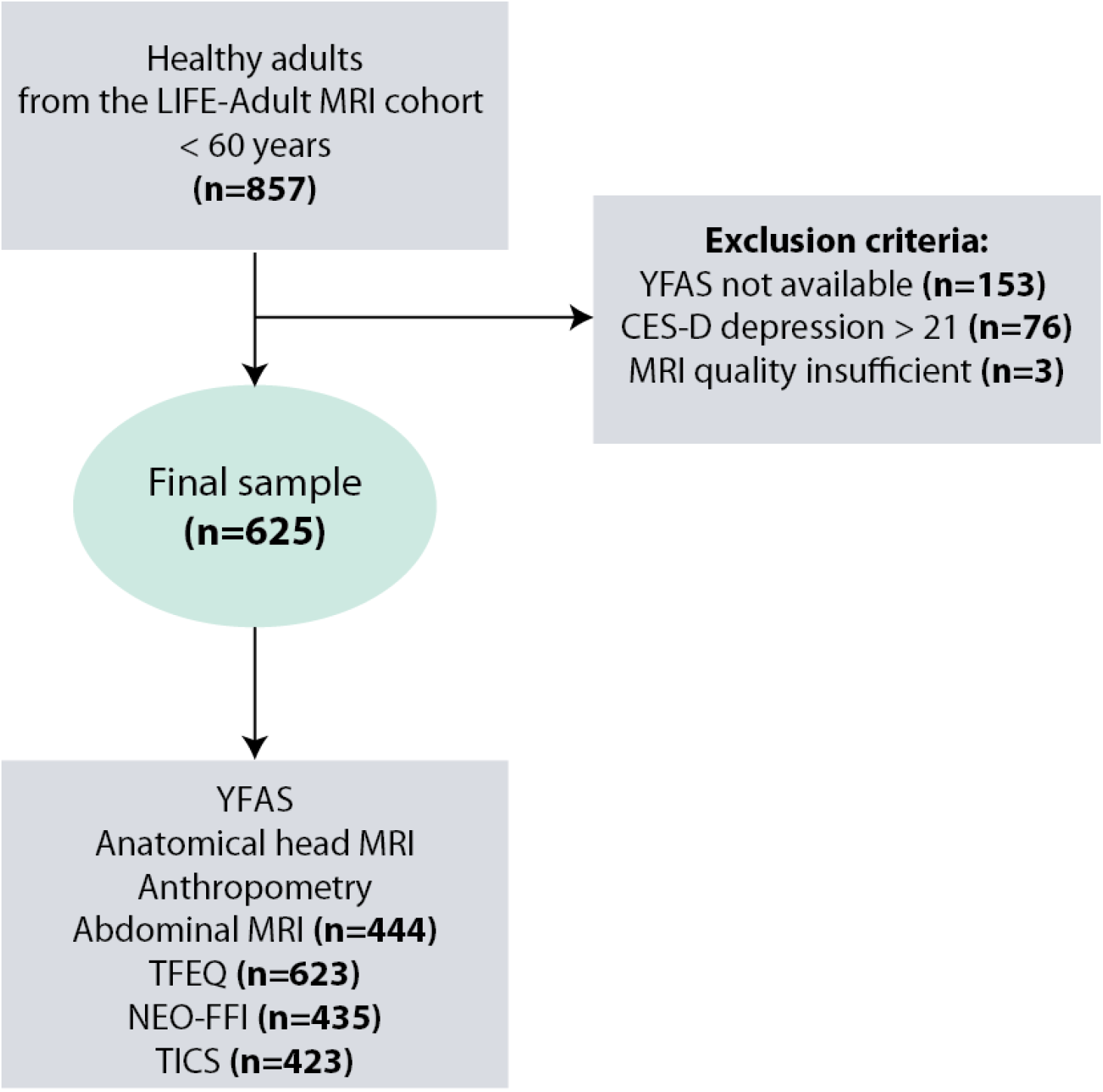
Flowchart showing the inclusion/exclusion criteria and variables of interest of the current study.

### Food addiction questionnaire

We measured food addiction using the Yale Food Addiction scale (YFAS) (Gearhardt et al., 2009). This questionnaire applied the DSM-IV criteria for substance dependence to eating behavior. The items of the scale have shown good internal consistency (Meule et al., 2012). We selected the continuous food addiction symptom score ranging from 0 to 7 as the primary variable of interest in our analyses as only 6% of the population meet criteria for manifest food addiction (Flint et al., 2014), and food addiction, like other psychopathological traits, might be better represented in a continuum (Kozak and Cuthbert, 2016).

### Head Magnetic Resonance Imaging

Anatomical T1 images were acquired using a 3 Tesla Siemens Verio MRI scanner (Siemens Healthcare, Erlangen, Germany) with a 3D MPRAGE protocol with the following parameters: inversion time, 900 ms; repetition time, 2300 ms; echo time, 2.98 ms; flip angle, 9°; field of view, 256 × 240 × 176 mm^3^; voxel size, 1 × 1 × 1 mm^3^.

Cortical thickness was estimated using Freesurfer’s standard pipeline recon-all (version 5.3.0) (Dale et al., 1999). Cortical thickness data was smoothed with a Gaussian kernel of 10 mm full-width at half maximum on the fsaverage template (Fischl and Dale, 2000). All images were visually checked for misplaced tissue boundaries and 56 images (∼10 %) were manually corrected. Three participants had to be excluded due to motion-related scan artifacts and segmentation errors.

Subcortical volumes were obtained from automated labeling (Fischl et al., 2002). Previous studies have emphasized the importance of the fronto-striatal circuits in addictive-like behaviors (Tomasi and Volkow, 2013; García-García et al., 2014; García-García et al., 2017). We therefore included the following structures: thalamus, caudate, putamen, pallidum, accumbens, and amygdala. For each subcortical structure, we extracted the volume for the right and left hemispheres separately and divided it by the total intracranial volume to account for head size. Two participants were identified as outliers in the subcortical analysis and were excluded due to enlarged ventricles and subcortical hyperintensities.

### Obesity-related variables

We used 4 anthropometric measures to characterize obesity (Pou et al., 2007; Amato et al., 2013): (i) body mass index (BMI), (ii) waist-to-hip ratio (WHR), (iii) subcutaneous adipose tissue divided by height (SCAT), and (iv) visceral adipose tissue divided by height (VAT).

BMI was used as a main measure to reflect obesity, since it is widely reported in studies in the field. WHR was recorded since it is considered to reflect visceral fat deposition (Lee et al., 2008). In addition, a subgroup of participants (n=444) underwent an abdominal T1-weighted MRI scan (Raschpichler et al., 2013). This acquisition covered the abdominal region starting 10 cm proximal and ending 10 cm distal from the umbilicus with a layer thickness of 1 cm. Sequence parameters can be found in the appendix. The subcutaneous and the visceral adipose tissues were quantified by means of semi-automated image segmentations (Raschpichler et al., 2013) and normalized by height (VanItallie et al., 1990).

### Eating behavior and personality questionnaires

We additionally characterized eating behavior by means of the Three Factor Eating Questionnaire (TFEQ) (Stunkard and Messick, 1985; Löffler et al., 2015b; Löffler et al., 2015a) which provides 3 measures of human eating behavior: (i) cognitive restraint of eating, (ii) disinhibited eating, and (iii) hunger. General personality traits were assessed with the short version of the NEO-FFI questionnaire, which provides measures for neuroticism, extraversion, openness, agreeableness, and conscientiousness (Körner et al., 2008).

Chronic stress was assessed with the screening scale of chronic stress (SSCS) from the Trierer Inventar zum Chronischen Stress (TICS) (Schulz et al., 2004). CES-D (Center for Epidemiological Studies-Depression) scores were used for exclusion (CES-D > 21) and to control for sub-threshold depressive symptoms (Radloff, 1977; Klinkman, 1997).

### Statistical analysis

For all statistical analyses, we used log-transformed values of the YFAS symptom score and the obesity-related variables (except WHR) to ensure normal distribution of regression residuals.

### Linear regression analysis of food addiction, obesity and personality measures

We used multiple regressions to explore associations between YFAS symptom score and the variables of interest (i) measures of obesity (ii) eating-behavior and personality, controlling for age, sex (with males as reference category) and BMI. Analyses were performed in R version 3.2.3.

### Cortical thickness analysis

We applied whole-brain vertex-wise analyses of cortical thickness and YFAS symptom score using Freesurfer’s mri_glmfit. In Model 1, we included YFAS symptom score, age and sex, in Model 2, we included BMI, age and sex and in Model 3 we included YFAS symptom score, BMI, age and sex.

Clusterwise correction for multiple comparisons based on pre-computed Monte-Carlo simulation was performed (cluster-forming threshold p = 0.0001, familywise error (FWE) corrected p<0.05). We applied the same statistical models in R (version 3.2.3) to analyze differences in subcortical volumes. Here, we accounted for multiple testing using Bonferroni’s correction (α_BF_=0.05/12= 0.004) for the 12 regions of interest (6 per hemisphere).

### Region of interest analysis using Bayes factor

Extending the whole-brain analysis, we aimed to weight the evidence for a contribution of food addiction, BMI or a combination of both to structural differences in selected regions of interest using Bayes factors.

Bayes factors (BF) quantify how likely the data are to occur under the assumption of one model compared to another (Goodman, 1999). More precisely, the Bayes factor BF_12_ is calculated as the ratio of the likelihood of the data given model 1 over the likelihood of the data given model 2. According to (Kass and Raftery, 1995), a BF_12_ between 3 and 20 indicates positive evidence for model 1 over model 2, a BF_12_ between 20 and 150 indicates strong, and a BF > 150 indicates very strong evidence for model 1 over model 2.

Here, we first compared Model 1 (including age, sex and YFAS symptom score) and Model 2 (including age, sex and BMI) to a null model including only age and sex. If this comparison yielded positive evidence for either BMI or YFAS symptom score (defined as a Bayes Factor > 3, (Kass and Raftery, 1995)), we additionally compared the models including YFAS symptom score and BMI alone (adjusted for age and sex, Model 1 vs Model 2), and the model including BMI along with YFAS symptom score to the model including BMI alone (adjusted for age and sex, Model 3 vs Model 2). To assess the collinearity of age, BMI and YFAS symptom score, we calculated the variance inflation factor (VIF) with the package “car” 2.1-3 in R.

We selected (orbito)frontal cortex and nucleus accumbens as ROIs, as structural differences in these regions were previously related to food addiction (Maayan et al., 2011) and obesity (Rapuano et al., 2017; García-García et al., 2018). We extracted individual cortical thickness values based on the Desikan-Killiany parcellation in Freesurfer for lateral OFC and medial OFC, rostral and superior frontal cortex on both hemispheres.

We used the package ‘BayesFactor’ version 0.9.12 in R version 3.2.3 to calculate and compare Bayes factors.

## RESULTS

### Sample characteristics

**Table 1.** Demographic characteristics and psychological measures of the sample (n=625, sex distribution: 344 men and 281 women).

|  |  |  |  |  |  |
| --- | --- | --- | --- | --- | --- |
|  | N | Minimum | Maximum | Mean | Std. Deviation |
| Age [years] | 625 | 20 | 59 | 40.6 | 10.8 |
| Food addiction symptom score (YFAS) | 625 | 0 | 5 | 1.4 | 0.9 |
| Depression scores (CES-D) | 625 | 0 | 21 | 8.4 | 4.6 |
| BMI [kg/m<sup>2</sup>] | 625 | 17.7 | 55.4 | 25.7 | 4.5 |
| WHR | 625 | 0.64 | 1.17 | 0.89 | 0.09 |
| VAT [mm<sup>3</sup>/cm] | 444 | 0.7 | 42.4 | 10.1 | 7.5 |
| SCAT [mm<sup>3</sup>/cm] | 444 | 1.36 | 88.8 | 21.6 | 12.1 |
| Cognitive restraint (TFEQ, 0 - 21) | 623 | 0 | 20 | 6.4 | 4.2 |
| Disinhibited eating (TFEQ, 0 - 16) | 623 | 0 | 15 | 4.4 | 2.7 |
| Hunger (TFEQ, 0 - 14) | 623 | 0 | 13 | 3.6 | 2.7 |
| Neuroticism (NEOFFI, 0 - 4) | 435 | 0 | 3.7 | 1.4 | 0.7 |
| Extraversion (NEOFFI, 0 - 4) | 435 | 0.8 | 3.8 | 2.4 | 0.5 |
| Openness (NEOFFI, 0 - 4) | 435 | 0.7 | 3.7 | 2.3 | 0.6 |
| Agreeableness (NEOFFI, 0 - 4) | 435 | 0.8 | 4.0 | 2.9 | 0.6 |
| Conscientiousness (NEOFFI, 0 - 4) | 435 | 1.5 | 4.0 | 3.1 | 0.5 |
| Screening scale of chronic stress (TICS, range 0 - 48) | 423 | 0 | 37 | 13.9 | 7.1 |

The items of the food addiction scale showed an acceptable internal consistency in the current sample of 625 healthy adults (Cronbach’s α= 0.78). YFAS symptom score was not significantly associated with age, sex or depression scores (all p > 0.05) (see Figure 2). 56 participants (8.9%) showed three or more symptoms of food addiction. Among those, eight participants (1.3 %) fulfilled the diagnosis criteria for food addiction and additionally reported clinical impairment or distress (Meule et al., 2012).

**Figure 2.**
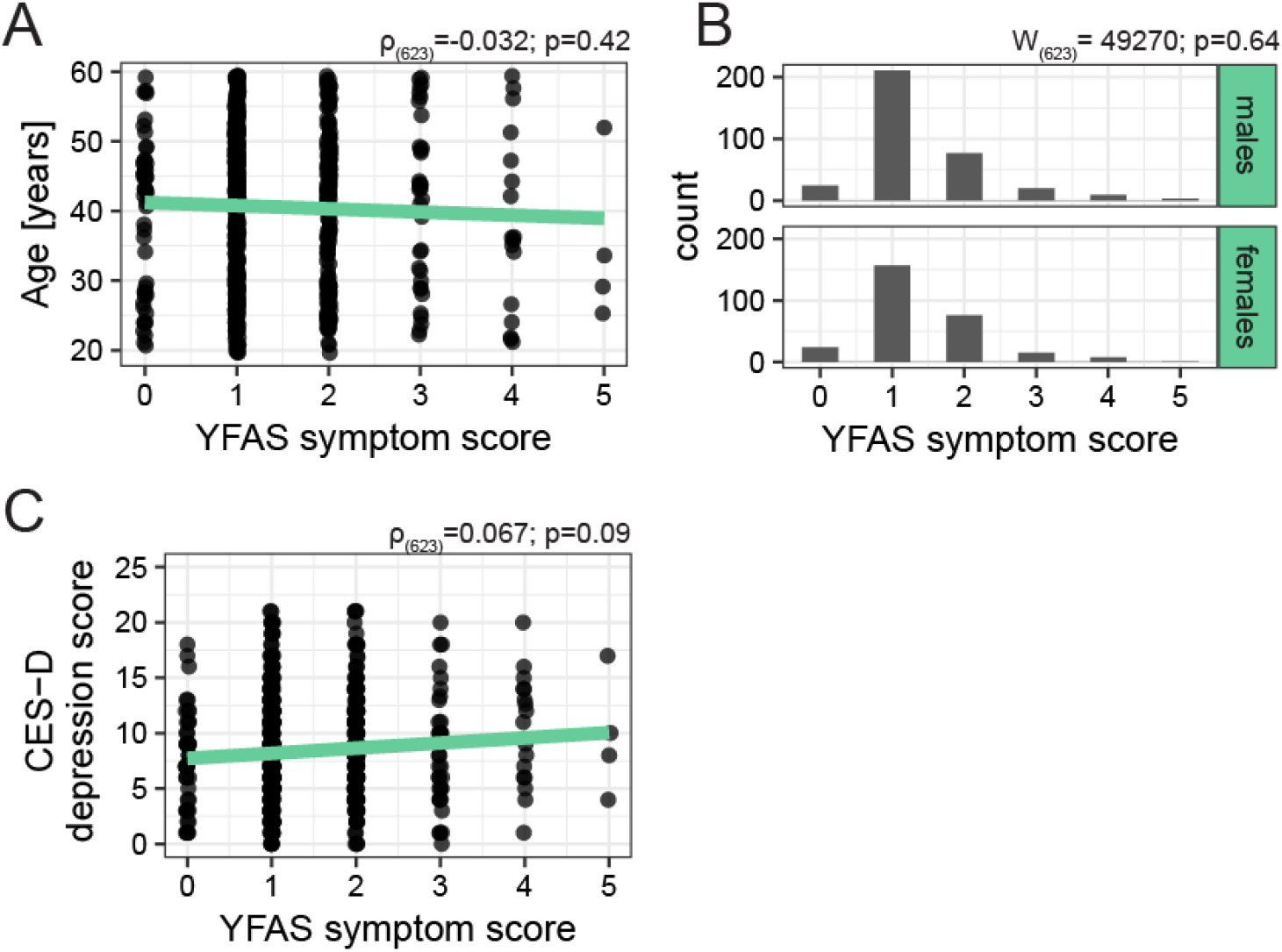
Relations between age (A), sex (B), CES-D depression score (C) and YFAS symptom scores in the investigated sample, visualized on the raw data. YFAS: Yale food addiction scale, CES-D: Center for Epidemiological Studies-Depression.

### Association of food addiction and obesity-related measurements

YFAS symptom score predicted BMI and WHR in the multiple regression analyses independent of age and sex. In the subgroup of participants with MRI-based measures of abdominal fat (n = 444), we also found that YFAS symptom score was positively associated with SCAT but not VAT (Figure 3; Table 2).

**Table 2.** Multiple regression analyses predicting different obesity-related measurements as a function of age, sex and YFAS symptom score

| | Adj. R<sup>2</sup> | B | C.I. | $\beta$ | P |
| --- | --- | --- | --- | --- | --- |
| **BMI** |  |  |  |  |  |
| Model | 0.22 |  |  |  | <0.001 |
| Age |  | 0.0027 | [0.0022 ,0.0032] | 0.41 | <0.001 |
| Sex |  | -0.03 | [-0.04,-0.02] | -0.18 | <0.001 |
| YFAS symptom score |  | 0.08 | [0.04, 0.11] | 0.16 | <0.001 |
| **WHR** |  |  |  |  |  |
| Model | 0.51 |  |  |  | <0.001 |
| Age |  | 0.0035 | [0.003, 0.0039] | 0.42 | <0.001 |
| Sex |  | -0.10 | [-0.11,- 0.09] | -0.56 | <0.001 |
| YFAS symptom score |  | 0.06 | [0.03, 0.09] | 0.10 | <0.001 |
| **SCAT** |  |  |  |  |  |
| Model | 0.23 |  |  |  | <0.001 |
| Age |  | 0.0082 | [0.0065, 0.01] | 0.39 | <0.001 |
| Sex |  | 0.13 | [0.09, 0.17] | 0.27 | <0.001 |
| YFAS symptom score |  | 0.22 | [0.10, 0.35] | 0.15 | <0.001 |
| **VAT** |  |  |  |  |  |
| Model | 0.54 |  |  |  | <0.001 |
| Age |  | 0.019 | [0.018,0.022] | 0.63 | <0.001 |
| Sex |  | -0.25 | [-0.29, -0.21] | -0.36 | <0.001 |
| YFAS symptom score |  | 0.09 | [-0.05, 0.24] | 0.041 | 0.19 |

**Figure 3.**
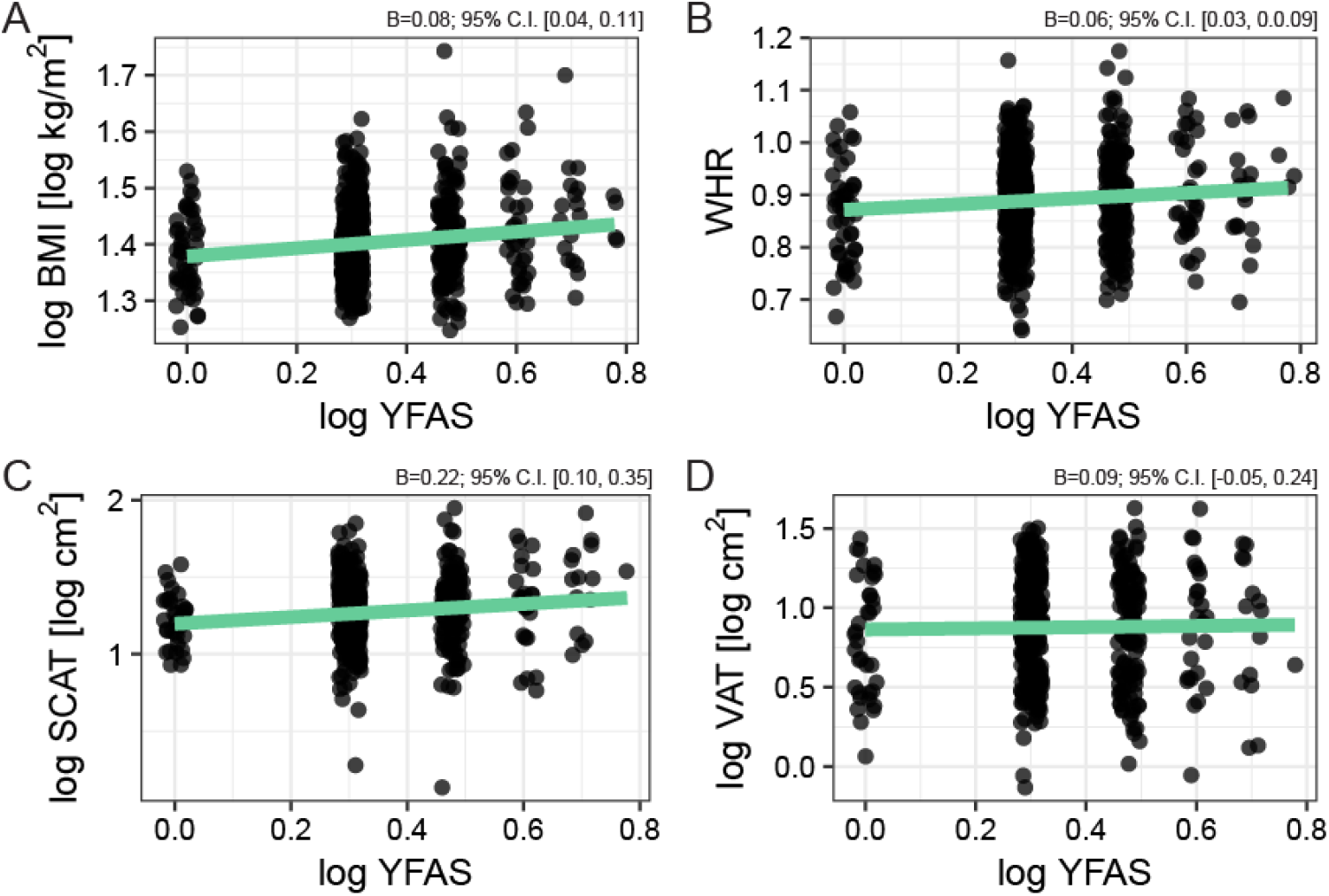
Associations YFAS symptom score (log-transformed) and (A) body mass index (BMI, log-transformed); (B) waist-to-hip ratio (WHR); (C) subcutanous adipose tissue (SCAT, log-transformed); and (D) visceral adipose tissue (VAT, log-transformed). The regression coefficient B and confidence interval (C.I.) are based on the multiple regression analysis accounting for age and sex.

### Relationship between food addiction, eating behavior and personality

In the multiple regression analyses, YFAS symptom scores explained variance in disinhibited eating (TFEQ), hunger (TFEQ), neuroticism (NEO-FFI) and chronic stress (TICS) after accounting for the effect of age, sex and BMI. In the four analyses, the directionality of the relation was positive (Figure 4; Table 3).

**Figure 4.**
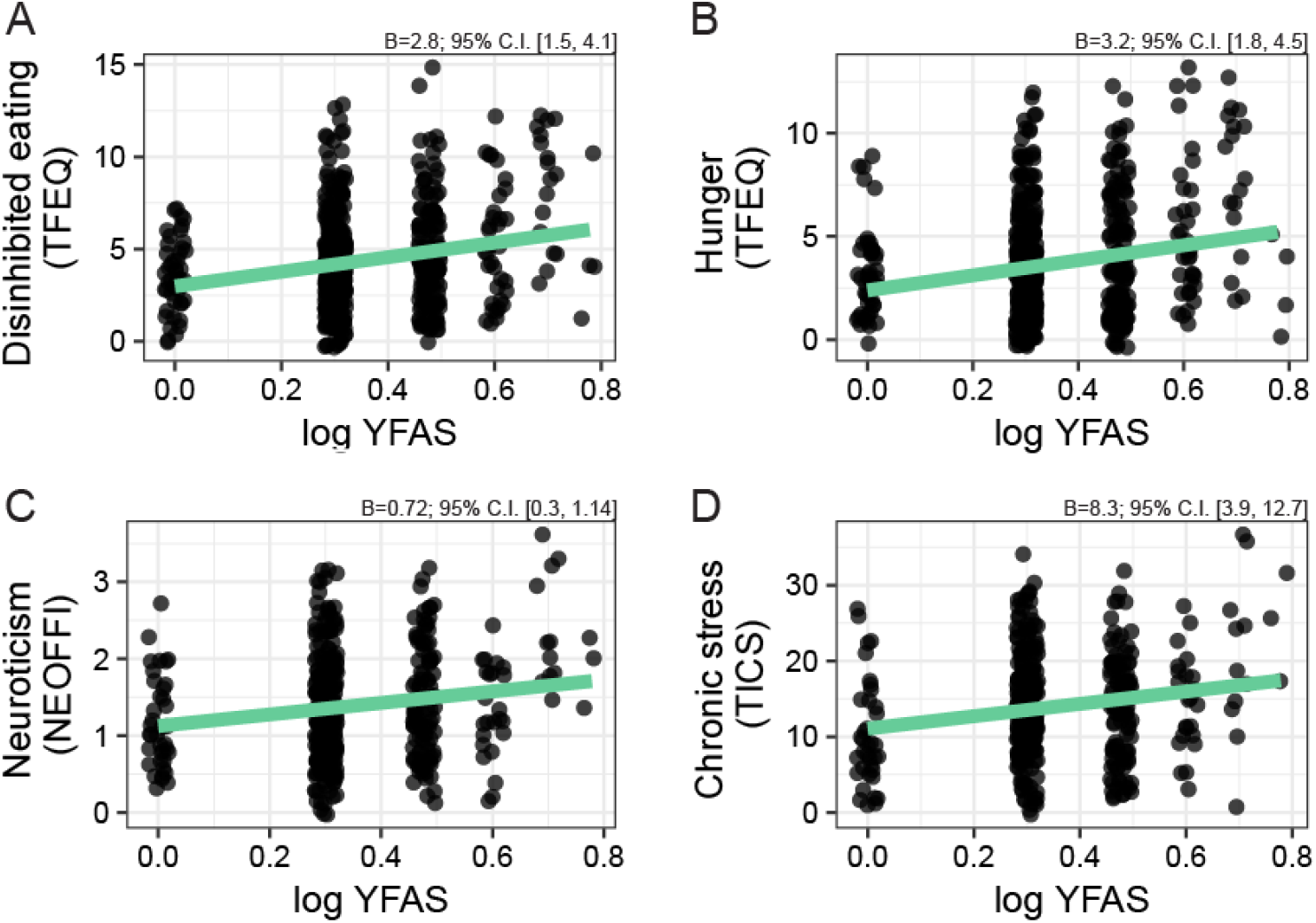
Significant associations between scores in YFAS symptom score (log-transformed) and (A) disinhibited eating; (B) hunger; (C) neuroticism; (D) chronic stress. The regression coefficient and confidence interval (C.I.) are based on the multiple regression analysis accounting for age, sex and BMI.

**Table 3.**
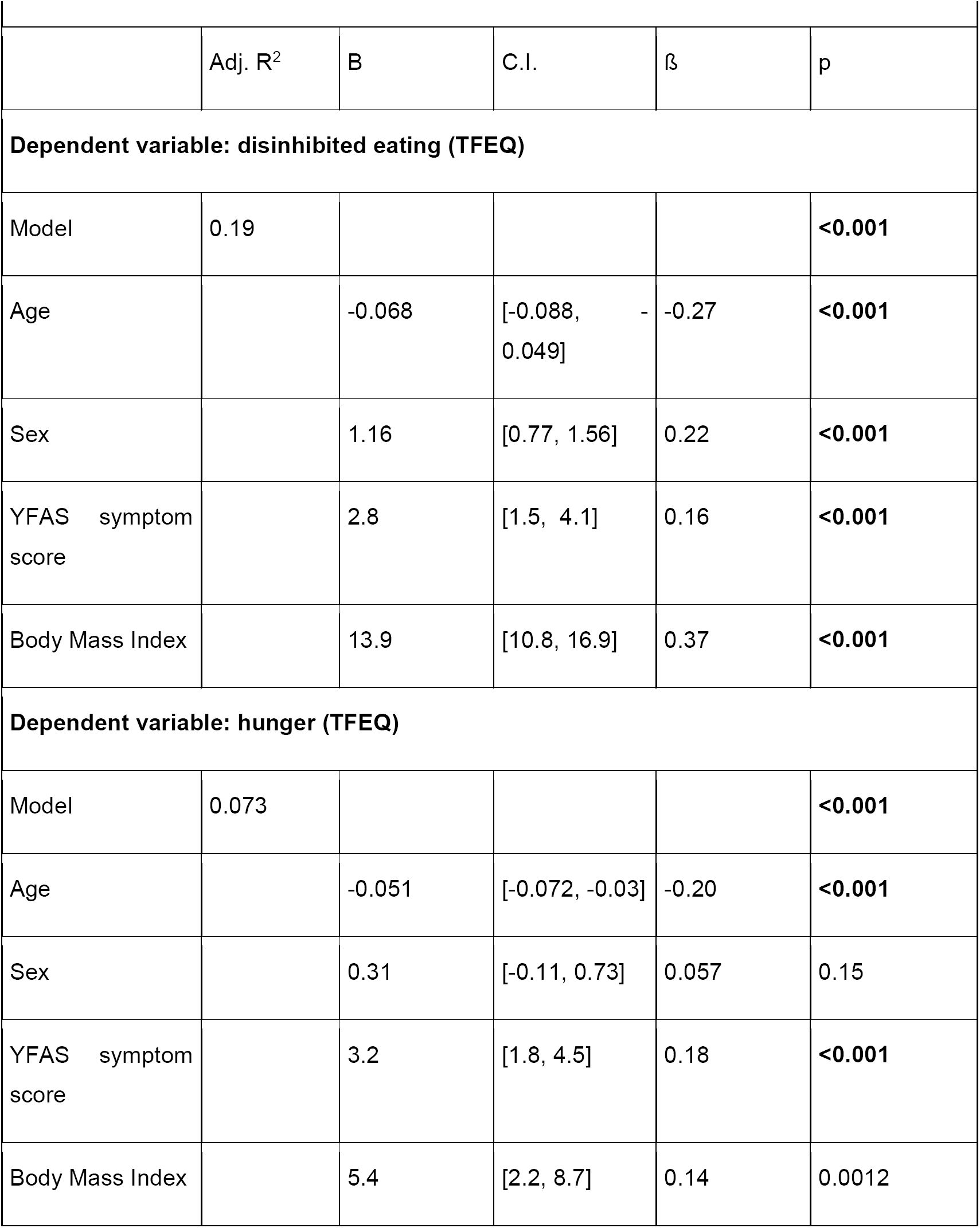
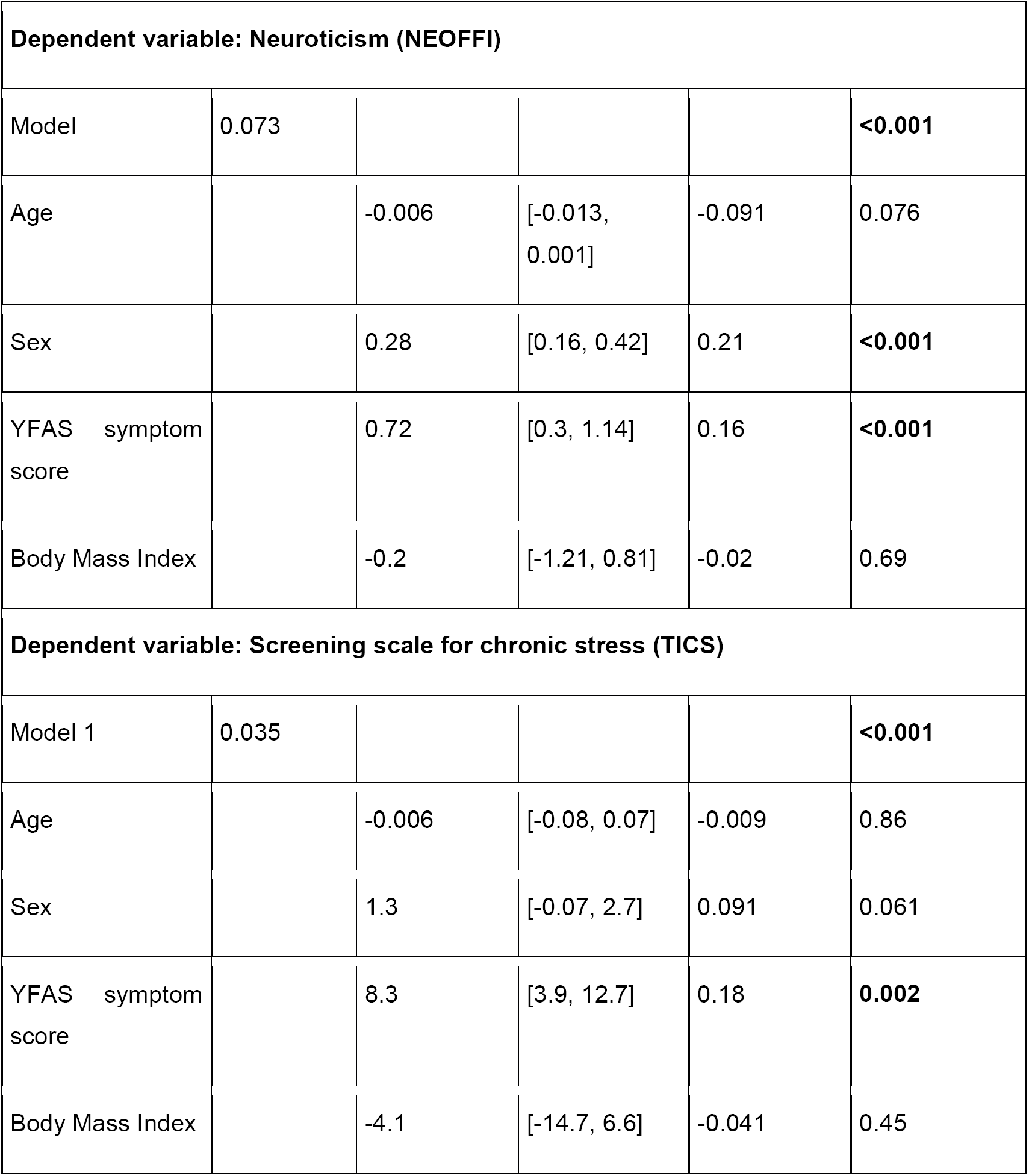
Multiple regression analyses predicting different eating behavior and personality measurements as a function of age, sex, YFAS symptom score and BMI

### Food addiction, obesity and cortical thickness

There was no statistically significant association of YFAS symptom score and cortical thickness with Model 1 (adjusting for age and sex) or Model 3 (additionally adjusting for BMI) (whole-brain FWE-corrected all p_cluster_ > 0.05).

Model 2 reveiled significant associations of BMI and cortical thickness adjusting for age and sex. Higher BMI was associated with cortical thinning in right lateral orbitofrontal cortex (OFC), rostral middle frontal, parahippocampal, isthmus cingulate and left lateral occipital cortex and middle and superior temporal cortex (see Table 4, Figure 5). When adjusting for YFAS symptom score (Model 3), all clusters except for the cluster in the isthmus cingulate remained significantly associated with BMI (Table 4; data not shown).

**Table 4.** Results of the cortical thickness regression analyses with YFAS symptom score (YFAS) and body mass index (BMI) for the right (rh) and left (lh) hemisphere

| Location | Cluster size (mm<sup>2</sup>) | MNI coordinates |  |  | P |
| --- | --- | --- | --- | --- | --- |
|  |  | X | Y | Z |  |
| **Model 1. YFAS** | None |  |  |  |  |
| **Model 2. BMI** |  |  |  |  |  |
| lh lateral occipital cortex | 555.47 | -43.9 | -77.8 | -11.2 | <0.001 |
|  | 176.22 | -41.4 | -77.5 | -1.6 | 0.0012 |
| lh middle temporal cortex | 415.44 | -48.4 | -5.8 | -30.6 | <0.001 |
|  | 82.58 | -50.5 | -18.2 | -15.9 | 0.032 |
| lh superior temporal cortex | 99.23 | -49.7 | -5.9 | -18.9 | 0.02 |
| rh parahippocampal cortex | 196.91 | 37.9 | -34.4 | -16.0 | <0.001 |
| rh rostral middle frontal cortex | 331.42 | 23.7 | 57.4 | -12.5 | 0.002 |
| rh lateral orbitofrontal cortex (OFC) | 428.59 | 27.3 | 18.8 | -10.5 | <0.001 |
| rh isthmuscingulate cortex | 128.30 | 6.1 | -45.4 | 30.7 | 0.007 |
| **Model 3.** |  |  |  |  |  |
| YFAS (<u>adjusted for BMI</u>) | none |  |  |  |  |
| BMI (<u>adjusted for YFAS</u>) |  |  |  |  |  |
| lh lateral occipital cortex | 451.3 | -43.9 | -78.1 | -11.0 | <0.001 |
|  | 95.1 | -41.4 | -77.5 | -1.6 | 0.02 |
| lh middle temporal cortex | 349.1 | -47.4 | -5.2 | -31.5 | <0.001 |
|  | 70.9 | -50.7 | -18.7 | -15.7 | 0.048 |
| lh superior temporal cortex | 114.6 | -49.9 | -5.3 | -18.8 | 0.01 |
| rh parahippocampal cortex | 88.26 | 37.8 | -33.4 | -16.4 | 0.02 |
| rh rostral middle frontal cortex | 155.04 | 23.7 | 57.4 | -12.5 | 0.0024 |
|  | 115.99 | 38.7 | 51.3 | 2.4 | 0.01 |
| rh insula/rh lateral OFC | 272.76 | 27.9 | 17.4 | -10.7 | <0.001 |
| **Model 1** we correlated YFAS symptom score and cortical thickness after controlling for age and sex. In **Model 2**, we tested the correlation between cortical thickness and BMI controlling for age, sex and in **Model 3** we included both YFAS symptom score and BMI controlling for age and sex. |  |  |  |  |  |

**Figure 5.**
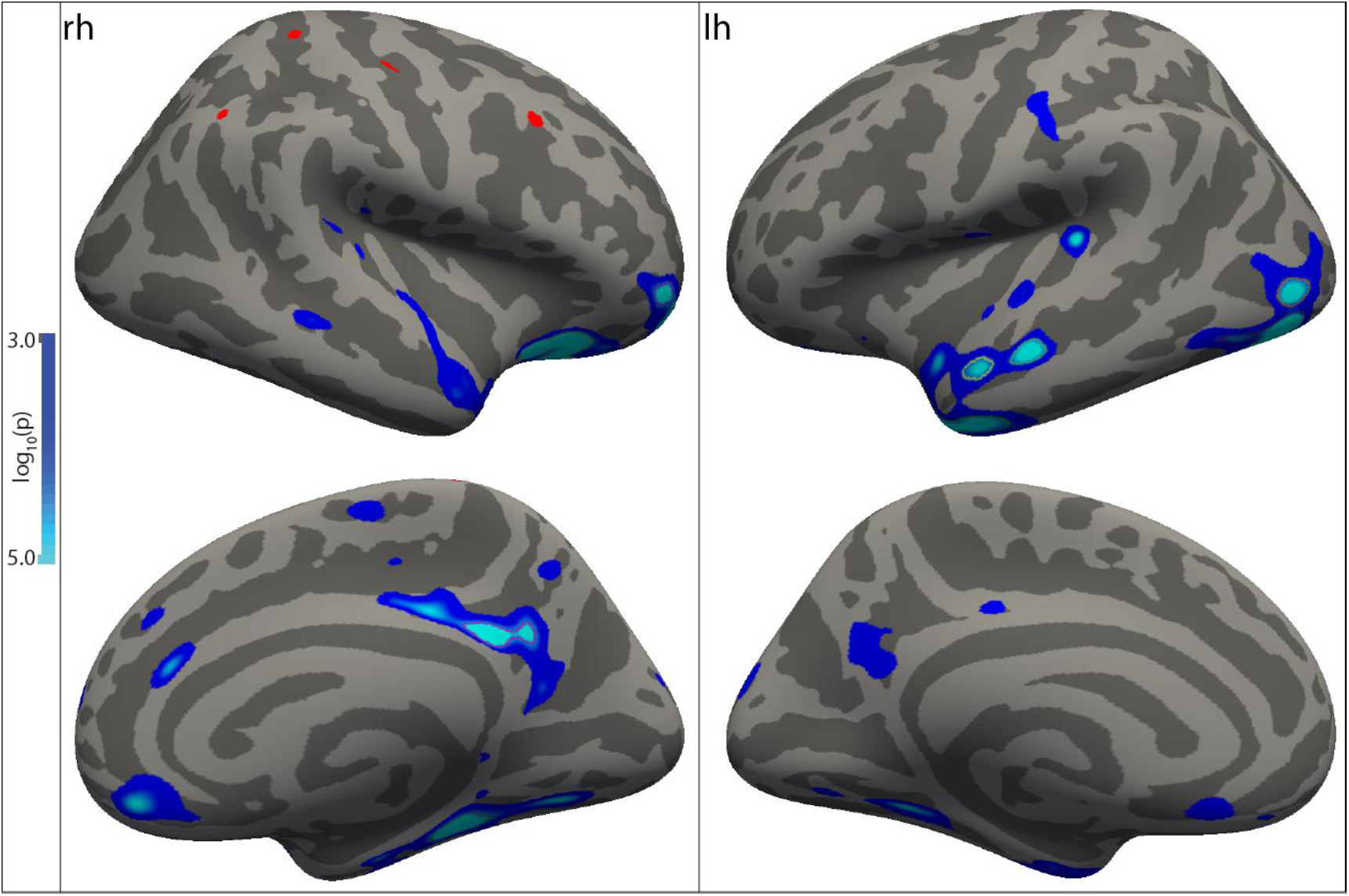
Higher BMI was associated with lower cortical thickness in the right lateral OFC, rostral middle frontal cortex, parahippocampal, isthmus cingulate and left lateral occipital, middle and superior temporal cortex after correcting for age and sex. The color bar represents the log10(p) values at each vertex projected onto the Freesurfer average brain.

### Food addiction, obesity and subcortical gray matter volumes

No significant correlations between YFAS scores and the subcortical volumes of interest emerged after age, sex (Model 1) and age,sex and BMI correction (Model 3) (all p > 0.004). BMI was significantly positively associated with increased left accumbens volume after correction for multiple comparisons (Model 2: standardized β = 0.15, p<0.001, Model 3: standardized β = 0.14, p=0.002). (for detailed results see supplementary table S1).

### Bayes model comparison

For the predefined regions of interest (right and left medial and lateral OFC, superior frontal and rostral middle frontal cortex, nucleus accumbens), we compared the evidence for models including age, sex and a) BMI b) YFAS symptom score and c) BMI and YFAS symptom score using Bayes factors (see Table 5).

**Table 5.** Results from the Bayesian model comparison between YFAS symptom score and BMI for gray matter volume/cortical thickness in predefined regions of interest. Bayes factors indicating strong evidence (BF > 20) are shown in bold.

|  | Bayes factors | YFAS vs null<br>(Model 1) | BMI vs null<br>(Model 2) | BMI vs YFAS<br>(Model 2 vs<br>Model 1) | BMI vs BMI +<br>YFAS<br>(Model 2 vs<br>Model 3) |
| --- | --- | --- | --- | --- | --- |
| rh | medial OFC | 5 | **266** | **52** | 0.86 |
|  | lateral OFC | **107** | **1997** | **18** | 0.09 |
|  | superior frontal | 0.14 | 0.59 |  |  |
|  | rostral middlefrontal | 0.28 | 2 |  |  |
|  | nucleus accumbens | 0.15 | 0.24 |  |  |
| lh | medial OFC | 0.33 | 0.38 |  |  |
|  | lateral OFC | 0.28 | 2.57 |  |  |
|  | superior frontal | 0.11 | 0.51 |  |  |
|  | rostral middle frontal | 0.11 | 0.2 |  |  |
|  | nucleus accumbens | **48** | 1.69 | **28** | 1.6 |

The variance inflation factors of age, BMI and YFAS were 1.22, 1.29 and 1.03, respectively.

For the right medial OFC, the analysis provided evidence in favor of a model including BMI, age and sex compared to a model including only age and sex (BF=266) or a model including YFAS symptom score, age and sex (BF=52). The data equally supported the model including BMI, YFAS symptom score, age and sex (BF=0.86).

In case of the right lateral OFC, there was very strong evidence for a model including BMI, age and sex (BF=1997). Strong evidence suggested a model including YFAS, age and sex (BF=107), however when comparing the age and sex-adjusted models with BMI and YFAS, the BMI model was preferred (BF=18). Positive evidence suggested including both YFAS and BMI (BF=11).

For right superior frontal, rostral middle frontal cortex and nucleus accumbens, the null hypothesis was preferred over age and sex-adjusted models including BMI or YFAS symptom score (all BF of comparison against null < 1/3).

Regarding the left hemisphere, no positive evidence for an association of BMI or YFAS symptom score and OFC or frontal cortical thickness was found (all BF of comparisons against null < 1/3). In line with the linear regression, a Bayes factor of 48 suggested strong evidence for an association of BMI and nucleus accumbens volume. Including both BMI and YFAS yielded a model that was equally likely (BF=1.16), and including an age-and sex-adjusted model with BMI was favored over a model including YFAS symptom score (BF=28).

## DISCUSSION

YFAS symptom score explained inter-individual variance in obesity measures, uncontrolled eating behavior and negative emotionality in a population-based cohort of 625 participants. In a whole brain approach, YFAS symptom score was not significantly associated with cortical thickness. In contrast, higher BMI was independently associated with reduced cortical thickness in right prefrontal, orbitofrontal and parahippocampal cortex and left temporal and occipital cortex. In addition, Bayes factor ROI analyses suggested a small contribution of YFAS symptom score to the cortical thickness of right lateral OFC, in addition to BMI.

### Neural correlates of food addiction and obesity

In this well-powered analysis (Pardoe et al., 2013), we found no statistically significant association of YFAS symptom score and cortical thickness on a whole-brain level. Similarly, subcortical volumes were not associated with YFAS symptom score. Regarding obesity, our analysis largely confirmed previous findings of BMI-associated reduced gray matter volume/cortical thickness. We replicated consistently reported lower gray matter volume in the right medial prefrontal cortex related to obesity, more specifically in its rostral subdivision and the orbitofrontal cortex (García-García et al., 2018). BMI was also associated with reduced cortical thickness in right parahippocampal, left temporal and lateral occipital regions and increased left accumbens volume (Veit et al., 2014; Walhovd et al., 2014; Rapuano et al., 2017). As we used Freesurfer cortical and subcortical analysis, we did not assess the previously reported relation of BMI and cerebellar gray matter volume (Kharabian Masouleh et al., 2016; García-García et al., 2018).

All findings related to BMI remained essentially unaltered after adjusting for YFAS symptom score. These findings indicate that addictive-like eating behavior does not account for the major part of structural differences related to obesity in the general population.

Accordingly, the model comparison using Bayes factors indicated that BMI rather than YFAS symptom score explained cortical thinning in frontal regions of the right hemisphere, and increased volume of the left nucleus accumbens. For the right lateral OFC, the data however also supported a model including YFAS symptom score alone, though this was 20 times less likely. Yet, positive evidence indicated that both BMI and YFAS symptom score might be associated with cortical thinning in this region. These - compared to the whole-brain analysis - somewhat conflicting findings should be interpreted cautiously. On the one hand, the evidence based on Bayes factors in the ROI analysis is moderate, making a definite conclusion about the association of YFAS and OFC cortical thickness difficult. On the other hand, in whole-brain, family-wise error corrected analyses, the larger effect size of BMI score and other factors such as age and sex compared to YFAS symptom, as well as the collinearity of BMI and YFAS symptom score, might have masked the small effect of food addiction. The OFC is a cardinal structure in our understanding of impulsive-and/or compulsive-related behavior (Zeeb et al., 2010; Burguière et al., 2013). Structural alterations in the OFC have been associated with impaired ability for goal-directed behavior (Reber et al., 2017) and impulsivity (Matsuo et al., 2009) and several studies reported diminished cortical thickness or GMV in the OFC of individuals with substance-related disorders (Durazzo et al., 2011; Ersche et al., 2012; Chye et al., 2017). More specific to overeating behaviors, reduced OFC gray matter volume and thickness have been related to less restrained eating (Su et al., 2017) and unhealthy food choice (Cohen et al., 2011), although one study reported larger OFC volume in binge-eating disorder patients compared to healthy controls (Schäfer et al., 2010). Structural alterations in OFC thickness could provoke, or fail to inhibit, impulsive and compulsive eating behavior, which would be partially captured by the YFAS symptom score.

Meanwhile, we find strong evidence for an association of BMI and lower cortical thickness in the OFC which might result from the adverse metabolic consequences of obesity (Cazettes et al., 2011; Dingess et al., 2017), and thus not be causally related to food addiction (Marqués-Iturria et al., 2013; Kharabian Masouleh et al., 2016; Thompson et al., 2017). Obesity, and especially visceral fat accumulation, is known to coincide with adverse metabolic responses, such as increased low-grade inflammation or progressive insulin resistance (Van Gaal et al., 2006), that enhance the vulnerability of the brain tissue (Corlier et al., 2018). Therefore, even though an association of reduced OFC thickness and addictive-like eating behavior seems plausible and is somewhat supported by the data, the current cross-sectional study provides evidence for a negative association of obesity and (orbito)frontal cortex thickness, largely independent of food addiction. Future studies investigating the neurobiological mechanisms underlying food addiction and obesity should therefore be longitudinal, include more sensitive assessments of addictive-like or uncontrolled eating behavior (Vainik et al., 2015) and investigate brain function, in addition to structure, related to these eating behaviors, for example using task-based functional MRI (Schag et al., 2013).

### Food addiction related to obesity and personality

Although the percentage of participants with high YFAS symptom score is low in general populations (Pedram et al., 2013), addictive-like behaviors might contribute to the high prevalence of obesity in modern societies (Schulte et al., 2016)(Hauck et al., 2017). A previous study, for instance, reported that participants tend to experience food addiction symptoms in relation to high-fat and sweet or savory foods (Markus et al., 2017). This result suggests that homeostatic mechanisms might be overwritten in favor of hedonic or habit-based eating. Such ingestive behavior might ultimately lead to weight gain. In line with these observations, we obtained modest but consistent positive correlations between food addiction and different measures of obesity. Further, our findings of linear relationships between YFAS symptom scores and obesity measures highlight that food addiction per se might rather be regarded as a dimensional trait which, like other psychopathological traits, might not fit traditional 1 versus 0 (“all or none”) diagnostic categories (Kozak and Cuthbert, 2016). Our results support a certain collinearity between obesity and food addiction (Pedram et al., 2013; Gearhardt et al., 2014; Pursey et al., 2016). Thus, taking into account individual tendencies for addictive-like eating behaviors might inform clinical approaches targeting trans-diagnostic symptoms of addictive behaviors, such as heightened impulsivity or compulsivity.

In the present study, higher YFAS symptom score was also associated with increased scores in disinhibited eating and hunger, two scales that reflect overeating and subjective food cravings, respectively. It is indeed possible that food addiction, disinhibited eating and hunger are partly overlapping scales, since they all do represent hyperphagia and loss of control over eating. In fact, some authors have suggested that different eating-related questionnaires tend to capture individual variations in “uncontrolled eating”, a dimension reflecting decreased self-control over eating (Vainik et al., 2015).

We additionally observed that YFAS symptom score was significant predictors of neuroticism, a personality trait linked with increased sensitivity to punishment and for negative emotionality (Costa and McCrae, 1985). This result is in line with previous findings suggesting that increases in depressive symptoms or emotional dysregulation significantly contribute to food addiction (Pivarunas and Conner, 2015; Chao et al., 2017; Markus et al., 2017). Again, the relationship between negative emotionality and food addiction could be reciprocal. A longitudinal study on a large cohort of female adolescents observed that depressive symptoms at baseline predicted the development of overeating behavior, while overeating behavior also predicted the onset of depressive symptoms (Skinner et al., 2012). Note that in the current analyses, we avoided confounding of potential manifest depression by excluding participants with CES-D scores > 21 and the majority of those were females. Notably, we did not observe sex differences in the YFAS symptom score, which might point to a similar prevalence of food addiction symptoms in both women and men. However, please note that we excluded participants with neurological and psychiatric disorder or symptoms and focused on a healthy cohort. In this rather homogenous sample, presumable sex differences in food addiction, as seen in many psychiatric disorders (Dohrenwend and Dohrenwend, 1976) might have been masked out.

### Methodological considerations

We would like to acknowledge three important limitations of the current study.

First, we assessed addictive-like eating behavior with the YFAS symptom score, which has a limited range (0-7) with relatively little variation in the current population. It might therefore lack sensitivity for inter-individual differences in impulsive-compulsive eating behavior (Vainik et al., 2015). Second, we collected no measurement for binge eating behavior. Food addiction presents some clinical overlap with binge eating disorder, since both conditions entail loss of control over food consumption along with continued use despite adverse consequences (Vainik et al., 2015; Kessler et al., 2016; Schulte et al., 2016). Third, we cannot conclude on the directionality of the associations due the cross-sectional nature of our study. Strengths of our study include a large, well-characterized population-based sample with high-resolution cranial MRI, several proxies of obesity including highly sensitive MRI-derived estimates of abdominal and subcutaneous fat, extensive questionnaires, and strict controls for possible confounding effects in the statistical analyses.

### Conclusions and Outlook

The present study shows no association of food addiction symptoms and cortical thickness in a whole brain analysis. We confirm previous findings of obesity-related cortical thinning in frontal, temporal and occipital brain regions. ROI-based Bayesian analysis suggests that BMI and YFAS symptom score might be independently negatively associated with right lateral OFC thickness.

Our findings indicate that symptoms of food addiction do not account for the major part of the structural differences commonly associated with obesity. Therefore, one plausible interpretation of our cross-sectional finding is that obesity-associated differences in brain morphology represent consequences of metabolic factors related to obesity, even in young to middle-aged healthy adults. Considering the overall low intensity of addictive-like eating behavior in our cohort, it is likely that subtle functional rather than gross structural differences in frontal areas drive these behaviors. Finally, it is also possible, that a vicious cycle relates structural differences, addictive-like eating behavior and obesity. Structural differences in frontal brain regions, induced by obesity-related factors, might further exacerbate impulsive and compulsive aspects of eating behavior, similar as in addictive disorders, and thereby lead to weight gain or reduced dieting success (Stevens et al.; DelParigi et al., 2007; Janssen et al., 2017). Thus, strategies aiming to reduce impulsive and compulsive eating behaviors might be beneficial in the treatment of obesity.

Appendix: 1 file with 1 figure, 1 table

## Acknowledgments

We would like to thank all participants and staff of the LIFE-Adult study, as well as Anja Dietrich for helpful discussions. This work was supported by grants of the European Union, the European Regional Development Fund, the Free State of Saxony within the framework of the excellence initiative, the LIFE–Leipzig Research Center for Civilization Diseases, University of Leipzig [project numbers: 713-241202, 14505/2470, 14575/2470] and by grants of the German Research Foundation, contract grant number CRC 1052 “Obesity mechanisms” Project A1, A. Villringer/M. Stumvoll, SCHR 774/5-1, M. L. Schroeter, and WI 3342/3-1, A.V. Witte, and by a Branco Weiss Fellowship, Society in Science to J. Sacher, and by a NARSAD Young Investigator Award by the Brain & Behavior Research Foundation to J. Sacher, and by the Max Planck Society.

